# Locally deployed AI agents for CRISPR guide discovery from outbreak reports

**DOI:** 10.64898/2026.10.20.760479

**Authors:** Timothy M. Schuler, Andrew S. R. Zobel, Gabriel A. Schuler

## Abstract

Translating outbreak reports into computational work supporting diagnostic development requires biological interpretation, literature review, data retrieval, and specialized analysis. We developed a domain-specific agent harness to coordinate a CRISPR guide-discovery pipeline using locally hosted open-weight models. The agent interprets requests, reviews literature, assembles inputs, and initiates analysis; the pipeline assesses candidate guides for researcher inspection. Evaluation with Gemma 4 31B, Nemotron 3.5 Super VL, and Qwen 3.8 27B on an NVIDIA DGX Spark demonstrated completion of the computational workflow. Disease and pathogen identification were consistent across runs, while gene selection, strain coverage, and candidate yield varied. Nemotron 3.5 Super VL had the shortest reported times to evaluation. Qwen 3.8 27B applied pipeline evaluation criteria more consistently, but required longer evaluation times. The runs demonstrated the utility of smaller language models when coupled with structured bioinformatics workflows. These findings support the feasibility of locally deployed agent coordination for diagnostic research as frontier models add additional restrictions on biological tasks.

## Introduction

Early recognition of an infectious-disease threat creates an opportunity to begin preparing a diagnostic response. The World Health Organization’s Epidemic Intelligence from Open Sources (EIOS) initiative supports detection and assessment of public-health threats by bringing together publicly available information for expert review [1]. A selected report can also motivate research into detection approaches relevant to the suspected pathogen and affected population.

CRISPR guide discovery requires consideration of target sequence diversity, compatibility with the selected CRISPR system, and potential recognition of background sequences. Existing methods, including ADAPT, have demonstrated automated diagnostic design that accounts for genomic variation and predicted activity [2]. Applying computational tools to an outbreak-related question also requires decisions about the target gene, genomic dataset, and biological background. These decisions connect information distributed across publications, sequence databases, and descriptions of the event.

Language model frameworks such as BioChatter connect natural-language interaction with biomedical information and tools [3]. Yoon et al. reported autonomous agents identifying a previously unrecognized reverse transcriptase system and pursuing follow-up analyses [4]. We examine a complementary application: locally hosted open-weight models coordinating computational work for diagnostic development. Our evaluation asks whether these models can carry an outbreak-related request through an existing bioinformatics pipeline and produce actionable results.

## Methods

### Guide discovery pipeline

The pipeline searches NCBI databases for genome assemblies representing the target organism, host, and additional biological background. NCBI provides access to genome sequences and associated metadata [5]. Target sequences from multiple strains are aligned using MAFFT and assessed against the selected CRISPR system’s requirements, including applicable protospacer adjacent motifs and guide architecture. The architecture includes 5′ or 3′ handles required by the configured CRISPR system. The pipeline searched for RNA targets for each run. The pipeline exports candidate spacers and includes any supplied guide handles in the folding analysis.

Candidate spacers are screened against the supplied host and background sequences to identify potential unintended sequence recognition. In the Gemma, Nemotron, and Qwen runs, background BLAST screening was configured with a minimum query coverage of 80% and a minimum sequence identity of 80%. This assessment is conditional on the reference collection used in each run. Only spacers that pass background screening proceed to folding analysis. Candidate guides are evaluated using RNAfold from the ViennaRNA package to predict secondary structures at different temperatures [6]. We use these predictions to inspect internal base pairing in the guide’s target-binding region as a structural indicator of potential accessibility. Results are presented through a web interface that visualizes the guides for researcher inspection.

### Agent harness and domain instructions

We used Hermes Agent, an open-source harness that provides the model-tool execution loop and session state [7]. We configured Hermes with domain-specific instructions and tooling and integrated it with the bioinformatics pipeline. These instructions supply workflow guidance; executable tools perform the requested operations. The contribution evaluated here is this domain-specific configuration and integration.

The agent interprets the request, reviews literature, selects a candidate target gene, assembles inputs, and invokes the pipeline. The pipeline performs sequence analysis and candidate assessment. Configuration and result artifact locations support inspection; successful execution is assessed from the available run records. Figure 1 summarizes this workflow.

**Figure 1.**
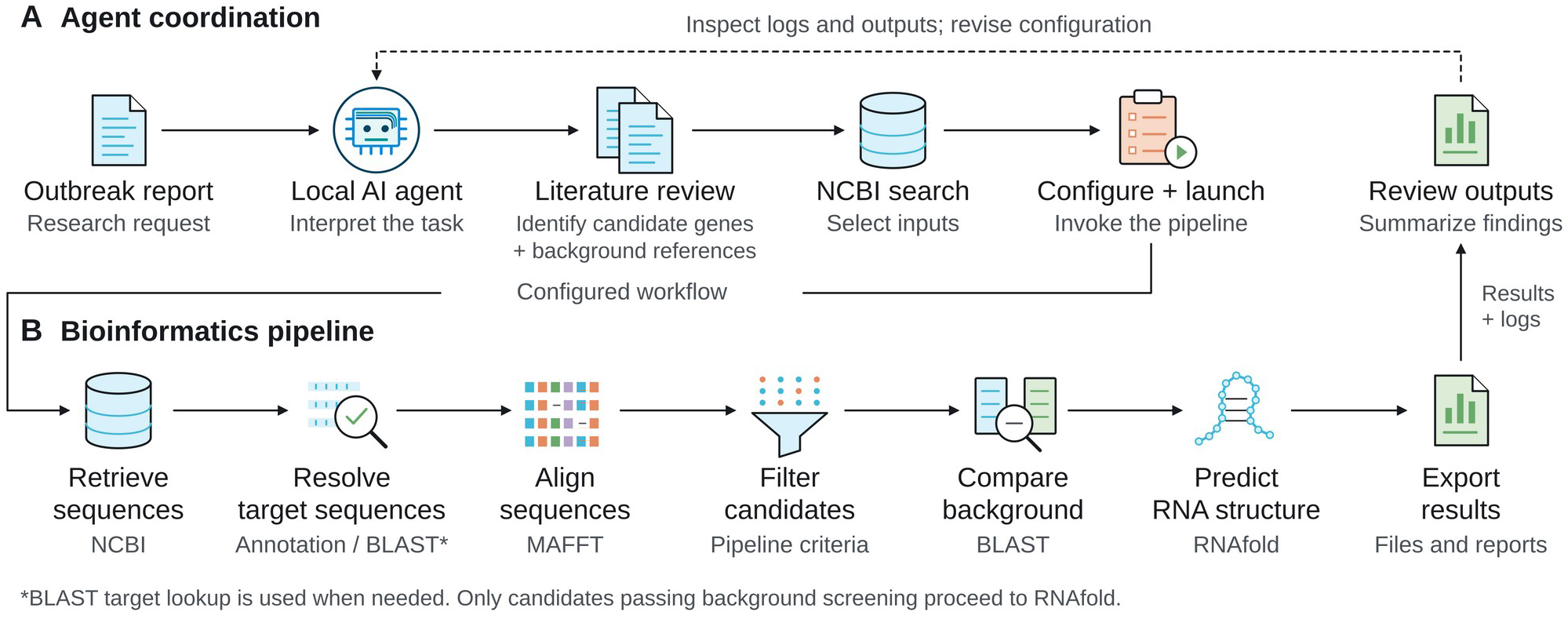
Agent coordination and bioinformatics workflow. (A) The local agent coordinates the research request, literature review, input selection, pipeline execution, and review of outputs. (B) The bioinformatics pipeline processes the selected inputs and exports results and logs. The dashed arrow indicates feedback to the agent.

### Local model evaluation

We evaluated three runs each for Gemma 4 31B, Nemotron 3.5 Super VL, and Qwen 3.8 27B. Nemotron 3.5 Super has 120 billion total parameters and 12 billion active parameters [8]. Runs used an outbreak-related article and a request to assist with computational work supporting diagnostic development. Across all runs, the agents correctly identified the disease as cholera from the WHO outbreak report [9]. As part of the literature review, the agents were prompted to focus on genes used as targets for PCR testing. The agents selected the gene and strain inputs for each run.

We recorded requested and resolved strain counts, selected gene, exported spacer count, reported strain coverage, artifact locations, and time to evaluation from the run summaries. Resolution refers to identification of the selected target sequence in an input strain; it is distinct from the coverage of an individual spacer. Coverage is expressed relative to the requested strains. We use the recorded session durations as the reported time to evaluation; for each model, we report the arithmetic mean, sample standard deviation, and range calculated from the original recorded durations. Session durations and means are presented to the nearest minute. All six Gemma and Nemotron runs proceeded without human intervention after the initial prompt. In all three Qwen runs, human involvement consisted of granting requested command-execution permissions; no changes were made to analytical decisions or workflow steps. Comparisons are descriptive.

### Computational environment

All model inference and bioinformatics analyses run on one NVIDIA DGX Spark with an NVIDIA GB10 Grace Blackwell Superchip, a 20-core Arm CPU, an integrated Blackwell GPU with 6,144 CUDA cores, and 128 GB of LPDDR5x coherent unified memory. NVIDIA specifies memory bandwidth of 273 GB/s [10].

## Results

All nine run summaries reported completed computational evaluations. Gemma 4 31B and Nemotron 3.5 Super VL each exported candidate spacers in two of three runs and returned no qualifying candidates in one run. Qwen 3.8 27B exported 13, 8, and 13 spacers across its three runs (Table 1). A completed evaluation could therefore yield either a candidate set or an empty result, depending on the analyzed inputs and configured criteria.

**Table 1.** Reported outcomes and time to evaluation.

| Model | Run | Strains Identified / resolved | Gene | Total Spacers Found | # Spacers after pipeline criteria | Reported Coverage Across Strains (%) | Time to evaluation |
| --- | --- | --- | --- | --- | --- | --- | --- |
| Gemma 4 31B | 1 | 2/2 | *toxR* | 5 | 5 | 100.00 | 28 min |
| Gemma 4 31B | 2 | 3/1 | *ctxA* | 13 | 0 | 33.33 | 26 min |
| Gemma 4 31B | 3 | 3/3 | *ctxA* | 13 | 13 | 100.00 | 32 min |
| Nemotron 3.5 Super VL | 1 | 3/2 | *ctxA* | 13 | 13 | 66.67 | 11 min |
| Nemotron 3.5 Super VL | 2 | 3/1 | *ctxA* | 13 | 0 | 33.33 | 10 min |
| Nemotron 3.5 Super VL | 3 | 3/3 | *hlyA* | 15 | 15 | 66.67 | 10 min |
| Qwen 3.8 27B | 1 | 3/3 | *ctxA* | 13 | 13 | 100.00 | 58 min |
| Qwen 3.8 27B | 2 | 3/3 | *tcpA* | 8 | 8 | 100.00 | 71 min |
| Qwen 3.8 27B | 3 | 3/3 | *ctxA* | 13 | 13 | 100.00 | 55 min |

Mean reported time to evaluation was approximately 10 min for Nemotron 3.5 Super VL (sample SD, 0.5 min; range, 10 to 11 min), 29 min for Gemma 4 31B (sample SD, 3.0 min; range, 26 to 32 min), and 61 min for Qwen 3.8 27B (sample SD, 8.5 min; range, 55 to 71 min). Nemotron evaluation times were shorter and more tightly clustered across these runs. These durations describe complete agent workflows with run-specific gene and strain choices.

All runs identified cholera and *Vibrio cholerae* from the outbreak report. Gemma and Nemotron each selected ctxA in two of three runs; the remaining selections were toxR for Gemma 4 31B and hlyA for Nemotron 3.5 Super VL. Gemma exported 5, 0, and 13 spacers across its three runs, with coverage ranging from 33.33% to 100%. Nemotron exported 13, 0, and 15 spacers, with coverage ranging from 33.33% to 66.67%. Qwen selected ctxA, tcpA, and ctxA, exporting 13, 8, and 13 spacers, respectively. All three Qwen runs resolved all three requested strains and reported 100% strain coverage, showing consistent completion and coverage despite variation in gene selection and candidate yield. Task interpretation was consistent across all three models, while gene selection, coverage, and candidate yield varied across runs.

The genes selected by the agents represent different aspects of *V. cholerae* biology and have distinct diagnostic interpretations. *ctxA* encodes cholera toxin subunit A and has been used as a toxin-gene target in molecular assays [11–14]. *tcpA* encodes the toxin-coregulated pilus subunit involved in intestinal colonization; its classical and El Tor variants have been used for biotype identification, with assay-dependent coverage [12–14]. *toxR* encodes a transcriptional regulator of cholera-toxin expression and has been used for species identification [12,13], whereas *hlyA* encodes hemolysin and was detected across the *V. cholerae* isolates examined in one study, including non-O1/non-O139 isolates [12]. Figure 2 summarizes the genomes selected across runs and the locations of candidate spacers in these four genes.

**Figure 2.**
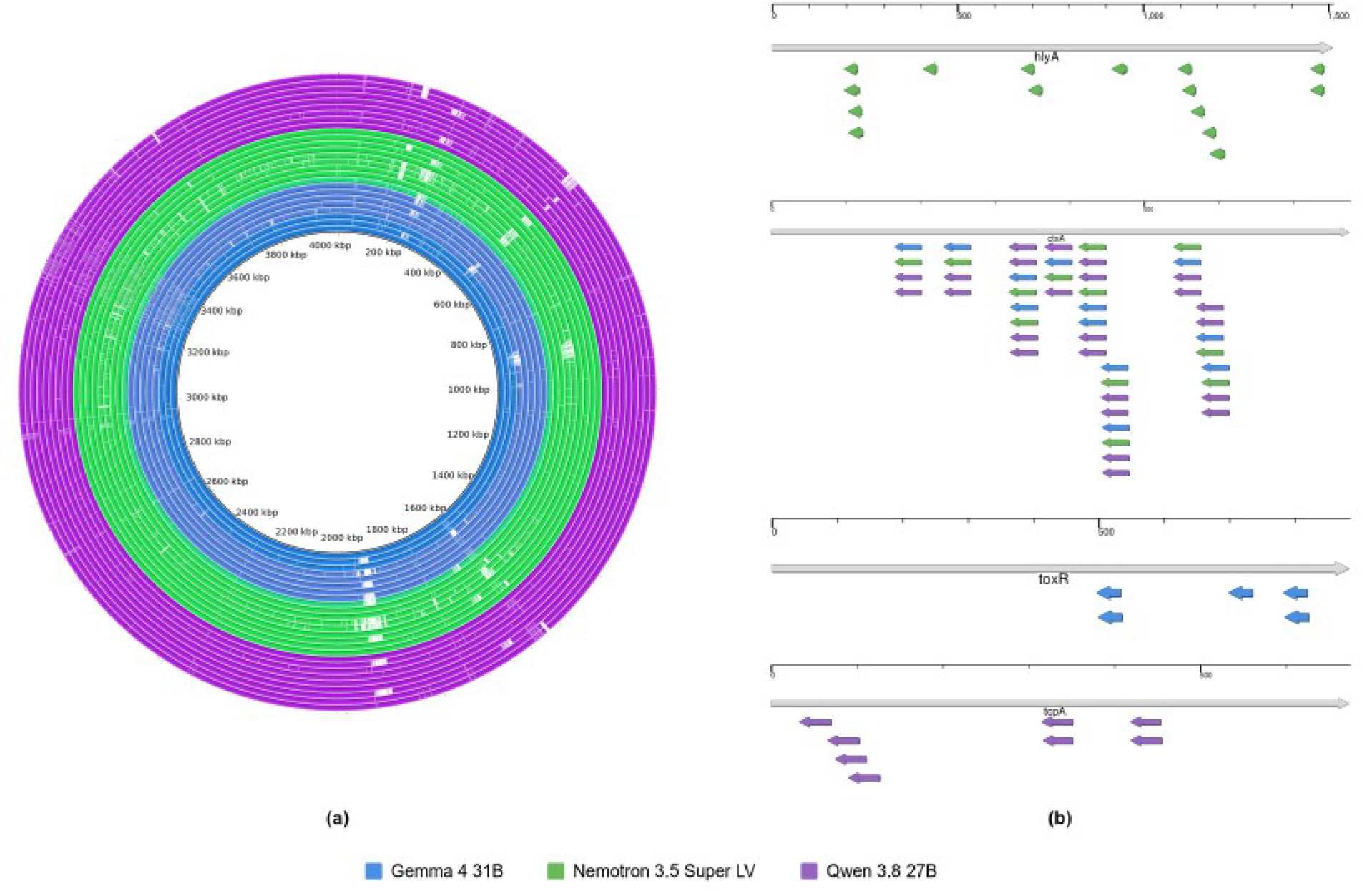
Genomes selected across runs and candidate-spacer locations. (a) Circular genome overview in which each concentric ring represents a genome selected for a given run. Coordinates are shown in kilobase pairs (kbp). (b) Candidate-spacer maps for *hlyA, ctxA, toxR*, and *tcpA*, from top to bottom. Gray arrows indicate the genes and their orientations; colored arrows show the positions and orientations of candidate spacers. Overlapping candidates are offset vertically for visibility. Each horizontal ruler shows nucleotide positions within the corresponding gene. Colors identify the model associated with each genome or candidate. Listed from innner to outer ring: Gemma 4 31B (blue, runs 1-3), Nemotron 3.5 Super LV (green, runs 1-3), and Qwen 3.8 27B (purple, runs 1-3).

The zero-spacer evaluations for Gemma and Nemotron each reported coverage of one of three requested strains (33.33%). Both returned an empty candidate set because this was below the pipeline’s configured requirement for guides to cover at least 50% of the selected strains. The two models therefore agreed on this type of evaluation outcome, although their selected assemblies differed. The reported counts and coverage summarize workflow outcomes; candidate-sequence agreement across runs was not assessed.

The Gemma and Nemotron summaries and the first Qwen summary described use of a human reference for background screening. The first Nemotron run reported 13 exported spacers and 12 folding scores, reflecting the restriction of folding analysis to spacers that passed background BLAST screening. Its third run reported 15 exported spacers and 15 folding scores. The first Qwen run also reported 13 exported spacers and 12 folding scores: 12 candidates were shortlisted for validation after background screening, while one was flagged for background review and did not proceed to folding. Its run summary reported successful completion of all 12 internal pipeline checks, with no warnings. The second Qwen run reported annotation-based extraction for all three strains, alignment with MAFFT, 639 enumerated windows, eight spacers passing all filters, no flagged background matches, and folding scores for all eight. The third Qwen run exported 13 spacers: 12 passed all filters, while one was flagged for a human background hit. The exported total therefore differs from the clean shortlist count.

## Discussion

Run summaries illustrate how the agents adapted the workflow and documented the sources, inputs, tools, and configuration changes used during execution. These accounts can help researchers review decisions and trace resources, with citations, logs, and artifacts providing the basis for verification. The evaluations show consistent identification of the outbreak disease and pathogen alongside variation in the computational results.

For these tests, we intentionally gave the agents control over analytical decisions and pipeline execution to evaluate autonomous workflow coordination. During all three Qwen runs, we approved requested command-execution permissions while the model retained control of the analytical workflow. The harness also supports additional safeguards and opportunities for researchers to review progress and discuss decisions with the agent [7].

Future agents could monitor news and public-health reports, including through EIOS, and proactively initiate computational work for diagnostic development and laboratory review. This direction is consistent with the WHO EIOS Strategy 2024–2026, which emphasizes AI-supported analysis, interoperability, and connections with broader public-health initiatives [15]. Our work explores a complementary application by linking outbreak reports to computational workflows supporting diagnostic research. Building on existing use of satellite observations for outbreak forecasting [16], extending these agents to social media, satellite imagery, and other sensor feeds could strengthen biosecurity through earlier threat recognition and more timely diagnostic preparedness.

This study complements AI-assisted biological discovery [4] with locally hosted open-weight agents. This work comes at a time when frontier-model developers are introducing safeguards and controlled-access pathways for biological research assistance. Anthropic describes fallback to a less capable model for certain biological requests [17], while OpenAI’s Rosalind Workbench requires organizational approval for advanced Research mode [18]. Anthropic’s Life Sciences Verification Program similarly makes access to more permissive biology safeguards conditional on institutional verification [19]. These developments make provider policies and eligibility requirements a practical consideration for laboratories adopting agentic bioinformatics assistance. We attempted to use GPT-6 Astra to run the pipeline, but biological-content restrictions prevented execution. Local agents could offer university laboratories and biotechnology companies an additional bioinformatics option under institutional governance, reducing reliance on provider access policies and potentially lowering costs associated with per-token billing for hosted models.

We plan to extend this evaluation with additional runs and a broader range of locally deployed models. Prior use of a gene in PCR-based detection alone does not establish its suitability for a CRISPR assay. Guide quality and background completeness remain unestablished; computational coverage does not establish diagnostic sensitivity or specificity. Experimental validation is necessary.

## Author declarations

Funding: No external funding was received. Competing interests: All authors are co-founders of NiagaraBio, Inc. a company working on CRISPR-based diagnostics. This work is associated with a provisional patent application.

